# Specific Regulation of *TCP genes* by miR319

**DOI:** 10.1101/747790

**Authors:** Javier F. Palatnik, Detlef Weigel

**Affiliations:** Instituto de Biología Molecular y Celular de Rosario (IBR), Rosario, Argentina; Max Planck Institute for Developmental Biology, Tübingen, Germany

## Abstract

MicroRNAs (miRNAs) are major regulators of gene expression in multicellular organisms. Many of the evolutionarily conserved miRNAs in plants are encoded by small gene families. The miR159/miR319 family has six members of similar sequences sharing 17 nucleotides in *Arabidopsis thaliana*. The members of this miRNA family regulate genes encoding TCP (<u>T</u>EOSINTE BRANCHED1, <u>C</u>YCLOIDEA and <u>P</u>CF1/2) and MYB transcription factors. However, despite their sequence similarities, genetic evidence indicates that miR159 and miR319 fulfil different roles *in vivo*. Here, we confirm previous findings showing that *TCP* genes are not targeted by miR159. Thus, specific small sequence differences between the miRNAs allow for the specific regulation of *TCP* transcription factors by miR319 miRNAs.

## Introduction

MiRNAs constitute a class of approximately 21 nucleotides long small RNAs that post-transcriptionally repress target genes in plants and animals. They are processed from longer precursors harboring a foldback structure with the miRNA embedded in one of its arms [reviewed in (Achkar et al., 2016; Yu et al., 2017)]. In plants, these precursors are processed in the nucleus by a complex formed by DICER-LIKE1 (DCL1) and accessory proteins such as HYPONASTIC LEAVES1 and SERRATE (Achkar et al., 2016; Yu et al., 2017). Mature miRNAs associate with ARGONAUTE (AGO) proteins, preferentially AGO1 in Arabidopsis [reviewed in (Ma and Zhang, 2018)]. The miRNAs guide AGO complexes to target RNAs by base complementarity, inhibiting their translation and/or guiding them to cleavage (Achkar et al., 2016; Yu et al., 2017). Plant miRNAs can recognize target RNAs with different degrees of complementarity, and the number and position of miRNA/target mismatches, the miRNA/target interaction energy and the context of the target sequence have been shown to affect miRNA-mediated gene regulation in plants (Allen et al., 2005; Li et al., 2014; Liu et al., 2014; Mallory et al., 2004; Schwab et al., 2005).

Many of the evolutionarily conserved miRNAs are encoded by small gene families, with the mature miRNAs being either identical or differing by a few nucleotides, but produced from the same precursor arm (Meyers et al., 2008). The Arabidopsis reference genome contains six genes encoding members of the miR159/miR319 family (Palatnik et al., 2007). All mature miRNAs are contained in the 3’ arm of their respective precursors, which are processed by four sequential cuts by DCL1 (Bologna et al., 2009). Phylogenetic studies have suggested a common origin of all genes encoding members of the miR159/miR319 family (Li et al., 2011).

MiRNAs of the miR159/miR319 family show high sequence complementarity to several transcripts encoding transcription factors of the TCP and MYB class. Experimental evidence, however, indicates that members of this miRNA family fulfill different functions *in vivo* (Achard et al., 2004; Allen et al., 2007; Nag et al., 2009; Palatnik et al., 2003; Palatnik et al., 2007). Overexpression of miR319a or miR319b induces strong changes in leaf morphogenesis and crinkled leaves, which phenocopies higher order *tcp* mutants (Bresso et al., 2018; Efroni et al., 2008; Koyama et al., 2007; Koyama et al., 2010; Schommer et al., 2008). On the other hand, overexpression of miR159a or miR159b has no obvious effects on leaf morphology (Achard et al., 2004; Allen et al., 2007; Palatnik et al., 2007), and instead, results in stamen defects, similar to those seen in *myb33 myb65* double mutants (Allen et al., 2007; Millar and Gubler, 2005).

Genetic evidence also highlights differences in the regulatory networks controlled by these miRNAs in Arabidopsis. The activation tagged mutant, *jaw-*D, is a miR319a overexpressor with crinkled leaves (Palatnik et al., 2003). A suppressor screen of *jaw-*D recovered mutations in the miR319-binding site of *TCP4* (Palatnik et al., 2007). Furthermore, a point mutation in the sequence of mature miR319a results in narrow petals, which are reverted to wild-type shape by a compensatory mutation in *TCP4* (Nag et al., 2009). On the other hand, *mir159a mir159b* double mutants have curled hyponastic leaves, similar to the phenotype of plants expressing a miR159-resistant *MYB33* transgene that has point mutations in the miRNA-binding site (Allen et al., 2007). Furthermore, in *mir159a mir159b myb33 myb65* quadruple mutants, in which two *MIR159* genes and two of its *MYB* targets are simultaneously inactivated, near normal leaf shape is recovered, demonstrating that miR159a and miR159b are required *in vivo* to control the expression of genes encoding MYB transcription factors (Allen et al., 2007).

Our previous work showed that expression levels as well as sequence differences between miR159 and miR319 are essential for the selection of the target genes *in vivo* (Palatnik et al., 2007). As we realized that a duplicated figure panel had been incorporated in Figure 4B of that publication, we have repeated the entire experiment, and replicated our previous findings. Here, we present our confirmatory results, showing that miR159 miRNAs do not affect *TCP4* mRNA or protein expression in transient *in vivo* assays.

## Results

We analyzed publicly available small RNA-sequencing data (https://mpss.danforthcenter.org/) (Nakano et al., 2006) and determined the most abundant miRNA generated for each precursor of the miR159/miR319 family (Figure 1). The six mature miRNAs are identical at 17 of the 21 nucleotides (Figure 1). Flexibility in the processing of their precursors leads to a proportion of the mature miRNAs having slightly different lengths, or sequences being shifted by one or two nucleotides with respect to the most abundant small RNA generated from each precursor.

**Figure 1.**
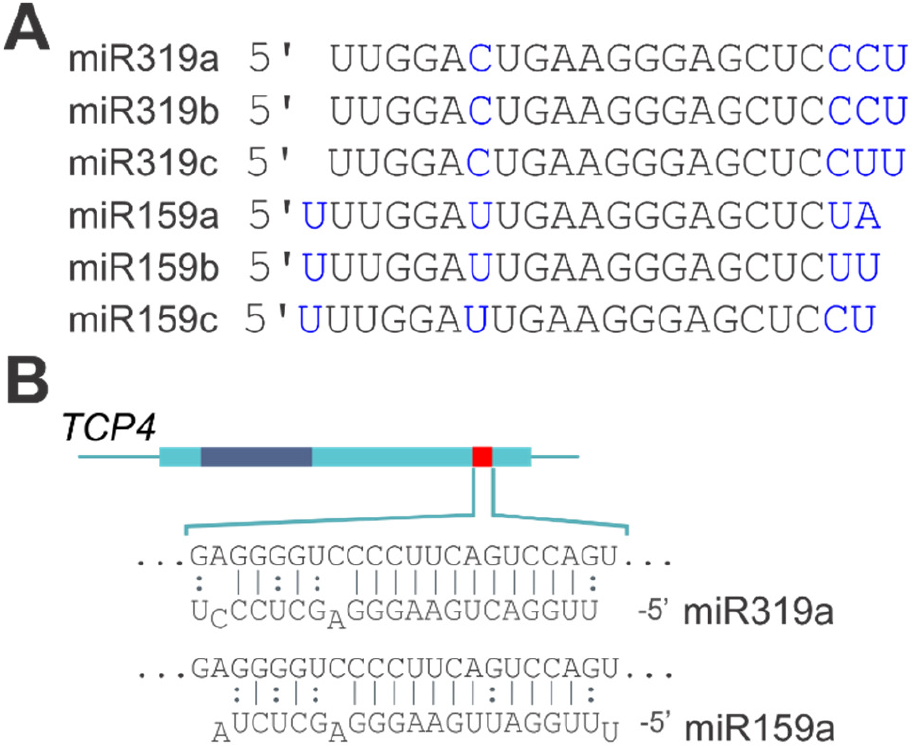
The miR159/miR319 family of miRNAs. (A) Sequences of the most abundant small RNA from each miRNA-encoding precursor. Blue letters highlight the differences. Note that miR159c accumulates to only very low levels. (B) Hybridization interface of *TCP4* with miR319a and miR159a. The miRNA complementary region in *TCP4* mRNA is highlighted as a red bar on top, and the region coding for the TCP DNA binding domain with a dark blue rectangle.

We analyzed with transient assays in *Nicotiana benthamiana* the effects of the different miRNAs on the accumulation of *TCP4* transcript and protein (Figure 2). Expressing a *TCP4:GFP* transgene in *N. benthamiana* leaves allowed us to detect *TCP4:GFP* transcript (Figure 2A) and GFP fluorescence (Figure 2B), despite the presence of substantial levels of endogenous miR159 in *N. benthamiana* leaves (Figure 2A). Co-expression of miR319a led to strong reduction of both, *TCP4:GFP* transcript and GFP fluorescence (Figures 2A and 2B). Even the weaker expression of miR319b (which is identical in sequence to miR319a) from a *35S:miR319b* transgene had a pronounced effect on *TCP4:GFP* RNA levels (Figure 2A). As a control, we used a version of *TCP4:GFP* with multiple mutations in the miRNA complementary motif (*mTCP4*), as described in Palatnik et al., 2003. *mTCP4:GFP* RNA levels appeared unchanged when miR319 was co-expressed, as expected for this miR319-insensitive version(Figure 2A).

**Figure 2.**
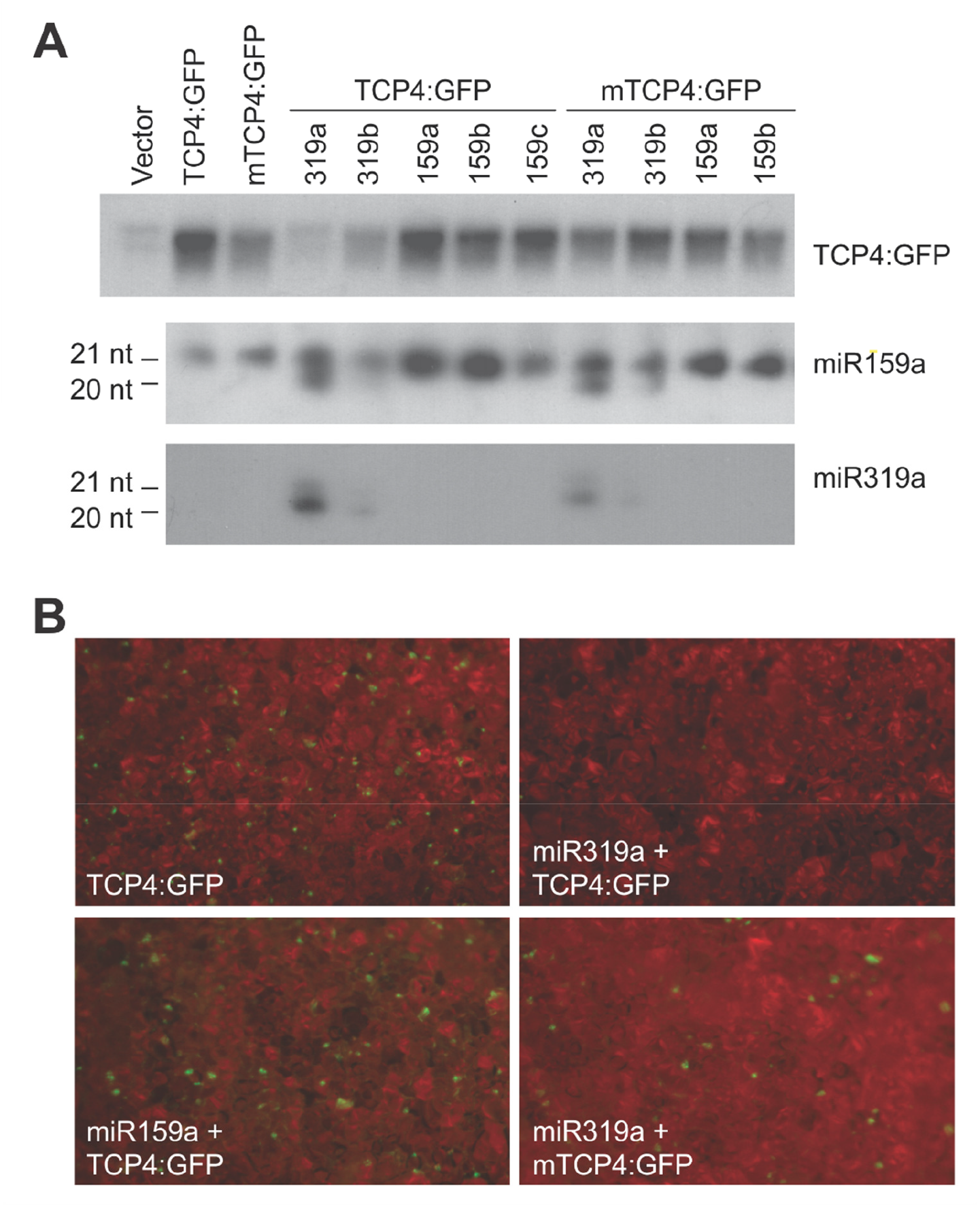
Specific regulation of *TCPs* by miR319. (A) Effect of different miRNAs on *TCP4:GFP* RNA levels in a *N. benthamiana* transient assay. (B) Effect of different miRNAs on TCP4:GFP fluorescence in *N. benthamiana* leaf epidermal cells.

Further increasing miR159 levels over the endogenous levels present in tobacco leaves by co-infiltrating *35S:miR159a* or *35S:miR159b*-encoding transgenes together with *TCP4:GFP* had no obvious effects on *TCP4* mRNA levels or GFP fluorescence (Figure 2A and 2B).

## Conclusions

Our results show that miR159 miRNAs do not regulate *TCP* mRNA or protein accumulation, and support conclusions that specific sequence differences between miR159 and miR319 underlie the different *in vivo* functions of the two miRNAs (Palatnik et al., 2007). Previous data showed that both the G-U wobble at position 7 of miR159, as well as the mismatches at the miRNA 3’ end prevent *TCP4* from being targeted by miR159 (Palatnik et al., 2007). The evidence is also consistent with data that modifications in the miR319-binding site of *TCP4* can compensate for changes in miR319 sequence (Nag et al., 2009) or levels (Palatnik et al., 2007). A similar situation is observed in the case of miR159 and *MYB* transcription factors, with *mir159a mir159b* mutant growth defects being suppressed by mutations in *MYB33* and *MYB65* (Allen et al., 2007).

## Material and Methods

The experimental approaches were performed as described previously (Palatnik et al., 2007).

## Acknowledgements

We thank Rebecca Schwab, Carla Schommer, Heike Wollmann, Ed Allen, Jim Carrington, Ramiro Rodriguez, Norman Warthmann and Jerôme Boisbouvier for comments on the manuscript and endorsing its content.

